# Functional dissection of neural circuitry using a genetic reporter for fMRI

**DOI:** 10.1101/2020.12.27.424403

**Authors:** Souparno Ghosh, Nan Li, Benjamin B. Bartelle, Tianshu Xie, Jade I. Daher, Urvashi D. Singh, Katherine Xie, Nicholas DiNapoli, Nicholas B. Evans, Kwanghun Chung, Alan Jasanoff

## Abstract

The complex connectivity of the mammalian brain underlies its function, but understanding how interconnected brain regions interact in neural processing remains a formidable challenge. Here we address this problem by introducing a genetic probe that permits selective functional imaging of neural circuit elements defined by their synaptic interrelationships throughout the brain. The probe is an engineered enzyme that transduces cytosolic calcium dynamics of probe-expressing cells into localized hemodynamic responses that can be selectively visualized by functional magnetic resonance imaging. Using a viral vector that undergoes retrograde transport, we apply the probe to characterize a brain-wide network of monosynaptic inputs to the striatum activated in a deep brain stimulation paradigm in rats. The results reveal engagement of surprisingly diverse projection sources and inform an integrated model of striatal function relevant to reward behavior and therapeutic neurostimulation approaches. Our work thus establishes a potent strategy for mechanistic analysis of distributed neural systems.

## INTRODUCTION

Optical measurements of signaling dynamics in well-defined neural circuit components are possible by combining invasive optical imaging methods with fluorescent probes like the genetically encoded calcium sensor GCaMP,^1^ but these approaches only operate over limited volumes. To permit measurements of circuit-level processing on a brain-wide scale, we sought to construct an analog of GCaMP that could be detected by noninvasive imaging methods like functional magnetic resonance imaging (fMRI),^2,3^ functional photoacoustic tomography,^4^ or functional ultrasound.^5^ These methods usually operate by monitoring blood flow changes evoked nonspecifically by neural population activity.^6^ We reasoned that a molecular probe designed to couple the intracellular signaling activity of genetically targeted cells to hemodynamic changes could form the basis of cell-specific functional neuroimaging, provided that output from the probe could be adequately distinguished from endogenous hemodynamic signatures (Fig. 1a). We refer to this concept as hemogenetic imaging, because it employs a genetic probe to produce a hemodynamic readout.

**Figure 1.**
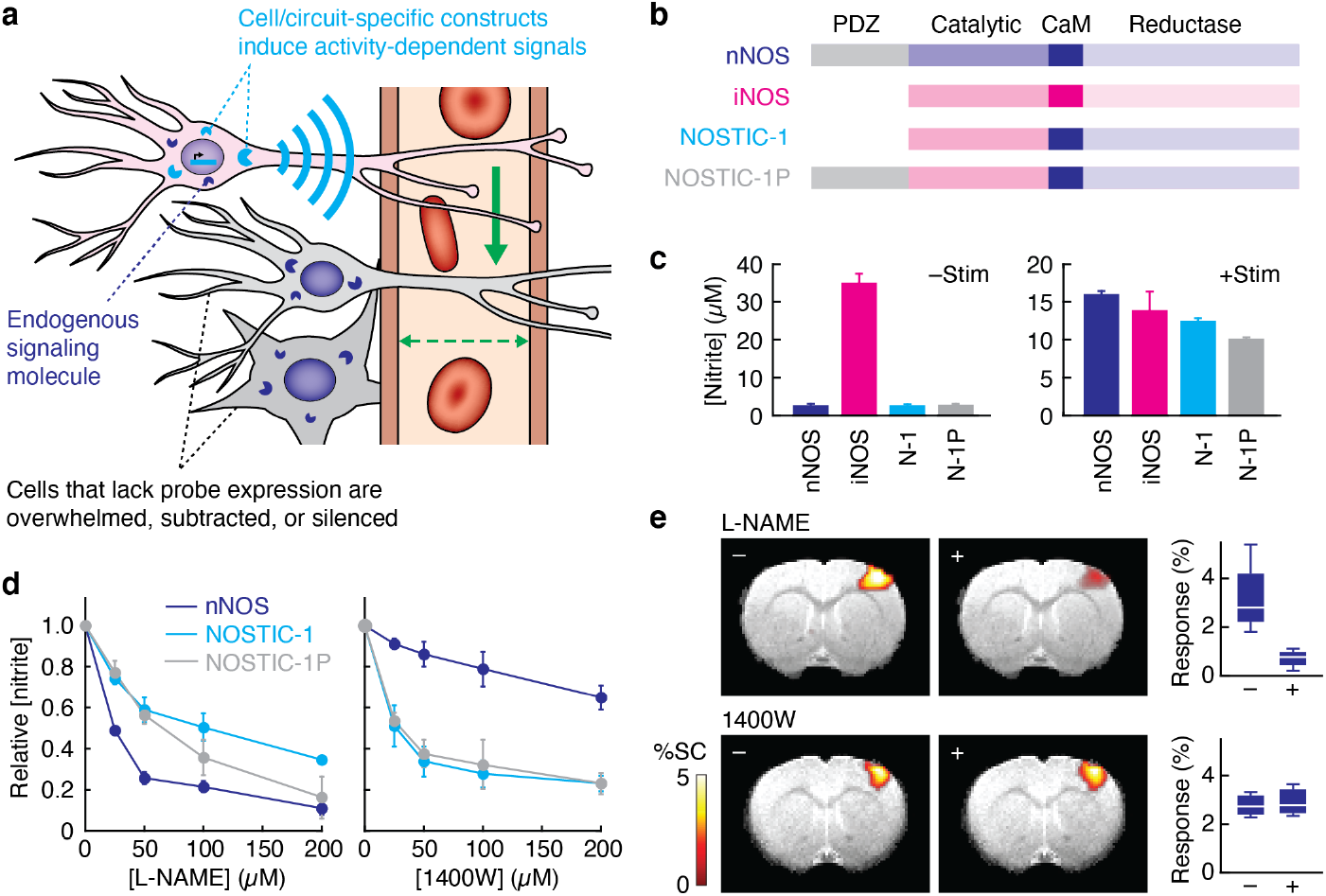
Principle of hemogenetic imaging with NOSTICs. **a,** Hemogenetic imaging involves detection of localized changes in blood flow or volume (green arrows) induced by genetically encoded activity reporters (cyan) analogous to fluorescent probes like GCaMP. Hemogenetic reporters signal to nearby blood vessels (cyan wireless symbol), inducing hemodynamic image contrast that can be detected noninvasively. Endogenous drivers of activity-dependent signals (dark blue) are suppressed, silenced, or subtracted. **b,** NOSTIC-1 and NOSTIC-1P reporters are calciumdependent chimeric enzymes that combine the catalytic domain of iNOS with calmodulin-binding (CaM) and reductase domains of nNOS; NOSTIC-1P also contains the synapse-targeting nNOS PDZ domain. **c,** Measurement of nitrite formation following NO release from cells expressing nNOS, iNOS, NOSTIC-1 (N-1), and NOSTIC-1P (N-1P) in absence (left) or presence (right) of stimulation using the Ca^2+^ ionophore A23187. **d,** Activity measurements titrated with the nNOS-selective inhibitor L-NAME (left) or iNOS-selective inhibitor 1400W (right). Error bars denote SEM of three measurements. **e,** Sensory-driven fMRI signals in rat cortex visualized with or without NOS inhibition using L-NAME (top, *n* = 4) or 1400W (bottom, *n* = 3). Images display percent signal change (%SC, color bar) superimposed on an anatomical scan. Box plots denote median (center line), first quartiles (boxes), and full range (whiskers) of fMRI responses observed in S1 forelimb region.

We chose a molecular basis for hemogenetic reporters by considering contributions to normal neurovascular coupling in mammalian brains. Although many signaling pathways have been implicated,^7^ a particular strong effect is mediated by neuronal nitric oxide synthase (nNOS). This enzyme catalyzes local formation of the cell-permeable gaseous vasodilator nitric oxide (NO) during neural signaling. Signaling activates nNOS via its association with calcium-bound calmodulin, the same protein domain that actuates fluorescent sensors like GCaMP.^8^ These considerations led us to design hemogenetic probes termed “nitric oxide synthases for targeting image contrast” (NOSTICs) by engineering an nNOS isoform that retains calcium sensitivity but that can be selectively manipulated by drugs that do not perturb intrinsic hemodynamic responses.

In this work, we sought to establish the principle of hemogenetic imaging with NOSTIC probes and begin to explore applications in neurobiology. Our results comprise protein engineering of the NOSTIC reporters, *in vitro* and *in vivo* validation studies, and functional analysis of neural circuitry in rodent brains using the approach. We show how hemogenetic fMRI provides brainwide information unavailable from other experimental methods and how this information can be combined with complementary data to yield insights into neural mechanisms.

## RESULTS

### Construction and validation of hemogenetic reporters

First-generation NOSTICs, called NOSTIC-1 and NOSTIC-1P, were created by replacing the catalytic domain of nNOS with corresponding residues from inducible nitric oxide synthase (iNOS), an immune-associated NOS variant that displays distinctive sensitivity to inhibitors (Fig. 1b).^9^ NOSTIC-1P further contains the synaptic targeting PDZ domain of nNOS,^10^ which NOSTIC-1 lacks. These probes therefore combine NOS activity with the possibility of cellular or subcellular spatial targeting.

Calcium dependence and pharmacological sensitivity of the NOSTIC probes were examined in cell culture. The calcium ionophore A23187 stimulates comparable levels of NO production by NOSTICs and nNOS expressed in HEK293F cells (Fig. 1c). The nNOS-selective inhibitor L-nitroarginine methyl ester (L-NAME) inhibits this effect with a half-maximal inhibitory dose (IC_50_) of 20 μM for nNOS, but 40-50 μM for the NOSTICs (Fig. 1d). In contrast, the iNOS-specific blocker 1400W inhibits NOSTICs with IC_50_ values of about 20 μM, with over tenfold lesser effect on nNOS (Fig. 1f). Importantly, the same two inhibitors produce sharply different effects on endogenous fMRI responses, as tested using a sensory stimulation paradigm in rats (Fig. 1e and Supplementary Fig. 1). While L-NAME injection reduces the cortical response to electrical forepaw stimulation by 2.41 ± 0.75% (significant with *t*-test *p* = 0.04, *n* = 4), 1400W produces virtually no change (–0.15 ± 0.05%, *t*-test *p* = 0.77, *n* = 3). These results thus demonstrate a basis for differential tracking of NOSTIC reporters by hemodynamic imaging.

To validate the probes *in vivo*, we examined the ability of NOSTIC-expressing cells to induce activity-dependent hemodynamic imaging signals following xenotransplantation into rat brains. HEK293 cells stably expressing NOSTIC-1 were implanted into the striatum and imaged using a 9.4 T magnetic resonance imaging (MRI) scanner during stimulation with A23187 (Fig. 2a and Supplementary Fig. 2). Calcium ionophore treatment induces robust blood oxygen level-dependent (BOLD) fMRI responses in the neighborhood of the cells (Fig. 2b). In contrast, control experiments performed in the presence of the NOSTIC inhibitor 1400W or using cells that express green fluorescent protein (GFP) in place of NOSTIC-1 result in negligible fMRI responses. The A23187-induced NOSTIC signal persists for ~10 min, likely reflecting slow washout of the ionophore (Fig. 2c), and results in a mean signal increase of 6.2 ± 1.4% (Fig. 2d) that is significantly greater than both controls (*t*-test *p* ≤ 0.001, *n* = 5). The full-width at half-height of the NOSTIC response averages 1.4 ± 0.1 mm, similar to the mean xenograft diameter of 1.8 ± 0.5 mm (Supplementary Fig. 3). This implies that the point-spread function width of NOSTIC-dependent fMRI signals is comparable to or less than the value of ~1 mm reported for intrinsic BOLD fMRI responses at 9.4 T.^11^ These experiments demonstrate that NOSTICs can selectively report the activation of probe-expressing cells in fMRI.

**Figure 2.**
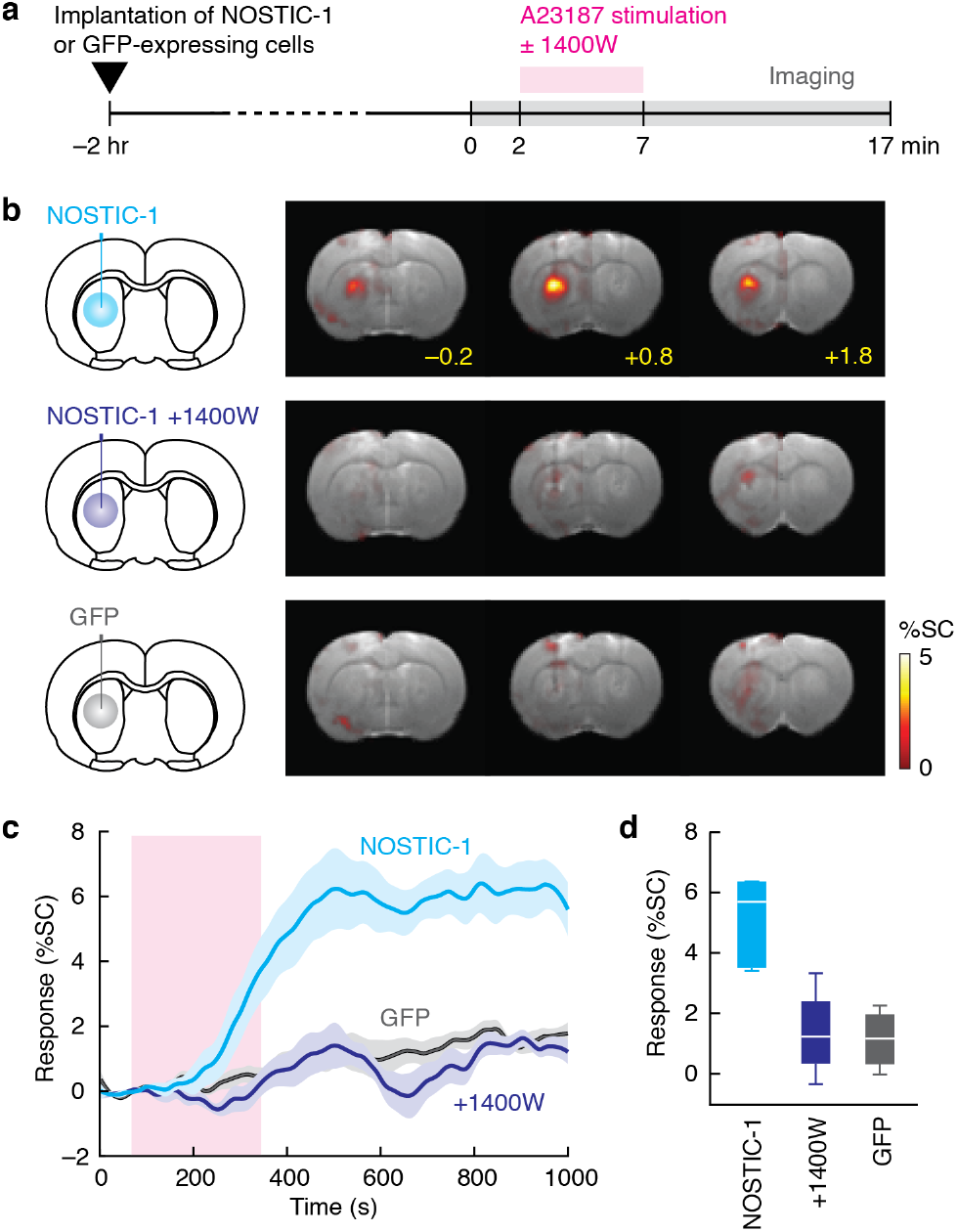
NOSTIC-1 reports activation of genetically-modified cells *in vivo*. **a**,Timeline of intracranial implantation and imaging of NOSTIC-expressing or control cells stimulated by infusion of A23187 in the absence or presence of the NOSTIC-1 inhibitor 1400W. **b,** BOLD MRI contrast changes induced by A23187-mediated stimulation: (top) NOSTIC-1-expressing cells without 1400W, (middle) NOSTIC-1 cells with 1400W, (bottom) GFP-expressing cells without 1400W. Each panel shows three coronal slices within 1 mm of the cell implantation site with a percent signal change (%SC) map overlaid on a representative anatomical scan. **c,** Mean time courses of MRI signals in ROIs centered around the cell implantation sites in **b** for animals implanted with NOSTIC-1-expressing cells in the absence (cyan) and presence (dark blue) of 1400W, and control GFP cells without 1400W (dark gray). Stimulation period denoted by light magenta box; shading denotes SEM of *n* = 6 (NOSTIC-1 −1400W), *n* = 4 (+1400W), or *n* = 4 (GFP). **d,** Average signal change amplitudes corresponding to the data in **c**. Box plots denote median (center line), first quartiles (boxes), and full range (whiskers).

### Functional imaging of targeted neural circuitry using NOSTIC-1

We next applied NOSTIC-1 to analyze neural circuit function in the striatum, a major hub for integration of sensory, motor, and affective input from remote brain areas, and a target of deep brain stimulation (DBS) therapies in the clinic. Understanding which connections provide active inputs during behaviorally relevant tasks or stimuli is essential to formulating circuit-level models of striatal function. Anatomical connections of the striatum have been studied by applying viral vectors that undergo axonal transport and express fluorescent proteins detectable in postmortem microscopy;^12–15^ functional assessment of the projections should therefore become possible by using similar vectors to drive expression of NOSTIC-1, followed by functional imaging. We used this approach to measure striatal input during electrical stimulation of the lateral hypothalamus (LH),^16^ a widely used addiction model in rodents and an exploratory DBS treatment for refractory obesity and depression in patients.^17,18^ A specific unanswered question in this context is whether striatal activation due to LH stimulation arises primarily from direct or indirect excitation of striatal afferents passing through this region,^19^ or whether alternative circuit-level mechanisms might help explain striatal responses and associated behavioral effects.

To label direct striatal afferents with NOSTIC-1, we constructed a herpes simplex virus (HSV) vector encoding NOSTIC-1 and mCherry (HSV-hEF1α-NOSTIC-IRES-mCherry). Similar vectors have been shown to produce persistent expression following retrograde transport from sites of injection in the rodent brain,^20^ implying that NOSTICs delivered this way should report the activity of neural populations that provide direct presynaptic input to striatal cells. After validating HSV-hEF1α-NOSTIC-1-IRES-mCherry in culture (Supplementary Fig. 4), we injected it into the anterior central caudate putamen (CPu) of six rats implanted with electrodes targeted to the medial forebrain bundle region of LH (Fig. 3a). Histological results confirmed that virally driven probe expression does not induce inflammatory markers (Supplementary Fig. 5). Three weeks after viral transduction, animals were sedated with medetomidine and imaged by BOLD fMRI during alternating blocks of LH stimulation and rest, first before and then after systemic treatment with the NOSTIC-1 inhibitor 1400W. These data permit identification of NOSTIC-specific signals based on the difference between fMRI responses observed in the absence *vs.* presence of the drug (Fig. 3a), reflecting monosynaptic functional input to CPu from throughout the brain during LH stimulation.

**Figure 3.**
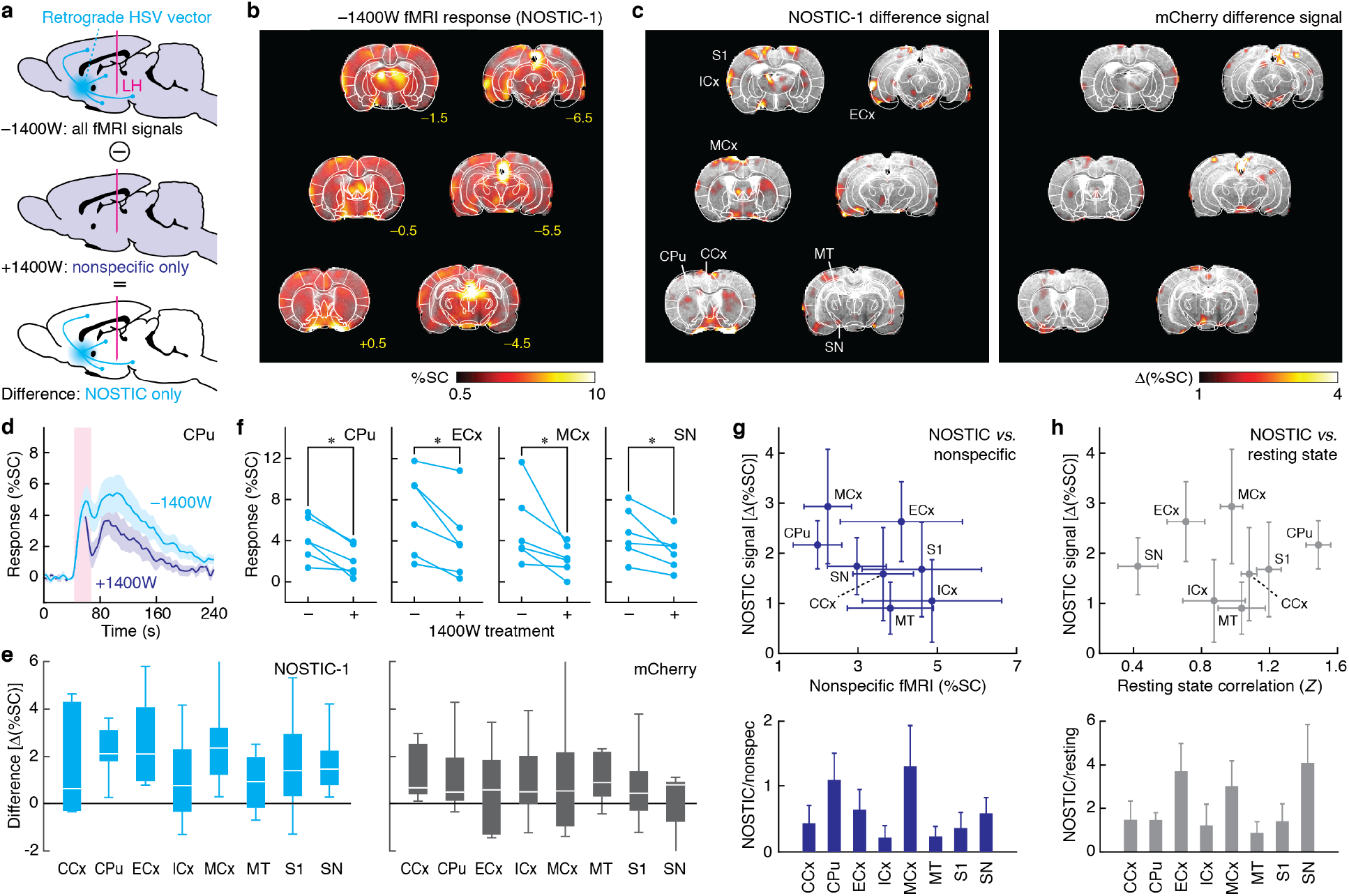
Hemogenetic functional imaging of striatal circuitry. **a,** Approach for hemogenetic analysis of striatal circuit function: An HSV vector delivers NOSTIC-1 to neurons presynaptic to a striatal injection site (cyan). Functional imaging during stimulation via an LH-targeted electrode (magenta) performed in absence (top) and presence (middle) of inhibition with 1400W, which suppresses NOSTIC-mediated fMRI signals but not endogenous nonspecific fMRI responses (purple). NOSTIC-specific responses to DBS are obtained by subtraction (bottom). **b,** Average fMRI responses to LH stimulation in the −1400W condition, among six animals infected with NOSTIC-encoding HSV. Significant responses with *F*-test *p* ≤ 0.01 shown. **c,** Mean amplitude of fMRI differences between −1400W and +1400W conditions observed for animals infected with NOSTIC-1 or mCherry-only viruses; select ROIs labeled. **d,** Average time courses of fMRI signals observed in CPu under each condition. Shading denotes SEM over six animals. **e,** Difference signal amplitudes for NOSTIC-1 (left) or mCherry control animals (right) in eight ROIs labeled in **c**(also see Supplementary Fig. 1). Box plots denote median (center line), first quartiles (boxes), and full range (whiskers). **f,** Absolute fMRI response amplitudes illustrating consistent 1400W-dependent responses in CPu, ECx, MCx, and SN. **g,** Comparison of estimated NOSTIC signals *vs.* nonspecific fMRI amplitudes recorded in the presence of 1400W (top). Ratios of NOSTIC to nonspecific signals shown at bottom. Error bars denote SEM (*n* = 6). **h,** Comparison of estimated NOSTIC signals (*n* = 6) *vs. Z*-transformed resting state correlation coefficients (*n* = 4) (top), with ratios shown at bottom. Error bars denote SEM.

A broad distribution of brain areas is activated both before and after 1400W treatment (Fig. 3b and Supplementary Fig. 1 and 6), but difference maps denoting activity specific to NOSTIC-expressing cells reveal a limited set of peaks (Fig. 3c). These peaks reflect 1400W-dependent scaling of the time courses of underlying fMRI responses, indicating that NOSTIC-dependent hemogenetic signals are temporally similar to endogenous BOLD responses (Fig. 3d). When averaged over eight anatomically based regions of interest (ROIs), difference amplitudes in several areas are substantially higher in NOSTIC-expressing animals than in rats transduced with the control vector HSV-hEF1α-mCherry (Fig. 3e and Supplementary Fig. 7). No significant ROI-level difference signals are observed in the controls (*t*-test *p* ≥ 0.08, *n* = 5), but 1400W-dependent fMRI signal differences from the NOSTIC animals are statistically significant (*t*-test *p* ≤ 0.05, *n* = 6) in CPu, entorhinal cortex (ECx), motor cortex (MCx), and substantia nigra (SN) (Fig. 3f). This suggests that NOSTIC-expressing cells in these regions provide the most consistent presynaptic input to the striatum during LH stimulation.

### Comparison of hemogenetic imaging with nonspecific fMRI

By comparing NOSTIC-mediated responses to intrinsic nonspecific fMRI signals from the same animals, recorded from the same animals in the presence of 1400W, the response profiles of neural populations with differing synaptic relationships to the striatum can be further distinguished (Fig. 3g). When normalized by nonspecific fMRI amplitudes, CPu and MCx display the strongest relative hemogenetic signals (about twofold greater than SN), suggesting that a particularly high proportion of stimulus-responsive cells in these two areas provides direct input to striatal HSV infection sites. In contrast, insular cortex (ICx) and medial thalamus (MT) stand out as strongly activated brain regions with the lowest mean NOSTIC-mediated responses, indicating that stimulus responses in these regions are elicited by striatal output or by pathways that are not directly related to striatal activity. These data imply that the DBS paradigm we studied engages neural populations far removed from the LH stimulation site; striatal function is shaped by local CPu circuitry and distal cortical projections, in addition to ascending input from midbrain neurons originating in SN.

We next considered whether hemogenetic measurements of circuit-level functional connectivity are similar to traditional “resting state” functional connectivity assessments derived from cross-correlation of regional fMRI time courses in the absence of tasks or stimuli.^21^ To explore this question, we acquired resting state fMRI data from four additional animals and computed *Z*-transformed correlation coefficients relating CPu time courses to each ROI (Fig. 3h). All eight ROIs display statistically significant *Z* scores (*t*-test *p* ≤ 0.001, *n* = 4). Interestingly, ECx, MCx, and SN all display relatively low values, despite identification of these regions as particularly robust striatal input sources using the NOSTIC probes; these areas exhibit the highest ratios of NOSTIC-mediated signal to resting state functional connectivity among all ROIs. The results thus suggest that all analyzed brain regions are functionally coupled to CPu activity, but that resting state analysis does not emphasize monosynaptic relationships engaged during LH-targeted stimulation.

### Integration of functional and anatomical measurements

Connectivity in neural systems is commonly studied using viral tracer-based methods, even though tracers alone do not provide functional information. We wondered to what extent NOSTIC-based functional measurements of striatal input are simply predictable from retrograde transport of striatally infused HSV vectors. Unbiased brain-wide histological analysis using SHIELD-based tissue clearing^22^ in animals infected by CPu injections of an mCherry-encoding HSV shows prominent fluorescence corresponding to cells in CPu, ECx, MCx, MT, and SN (Fig. 4a); additional features are observable in individual tissue slices (Fig. 4b). Comparison of HSV marker visualization with activity-dependent *c*-Fos protein expression following treatment similar to the experiments of Fig. 3b-g shows that CPu, ECx, MCx, and SN all contain co-stained cells, consistent with the NOSTIC-based reports (Fig. 4c). In MT, however, *c*-Fos-positive cells are distinct from those labeled with the virus (Fig. 4d). Quantification of mCherry fluorescence levels across regions of interest in the histological slices of Fig. 4b shows that the most highly HSV-labeled areas differ from those with the most robust NOSTIC-dependent activity signals in the experiments of Fig. 3 (Fig. 4e). MT stands out as a highly mCherry-labeled region which does not provide presynaptic input to the striatum during LH stimulation, whereas MCx is only modestly labeled by the virus, despite showing strong striatal presynaptic input from those cells that are labeled. These results therefore dissociate stimulus-driven circuit function from the anatomy of striatal afferents.

**Figure 4.**
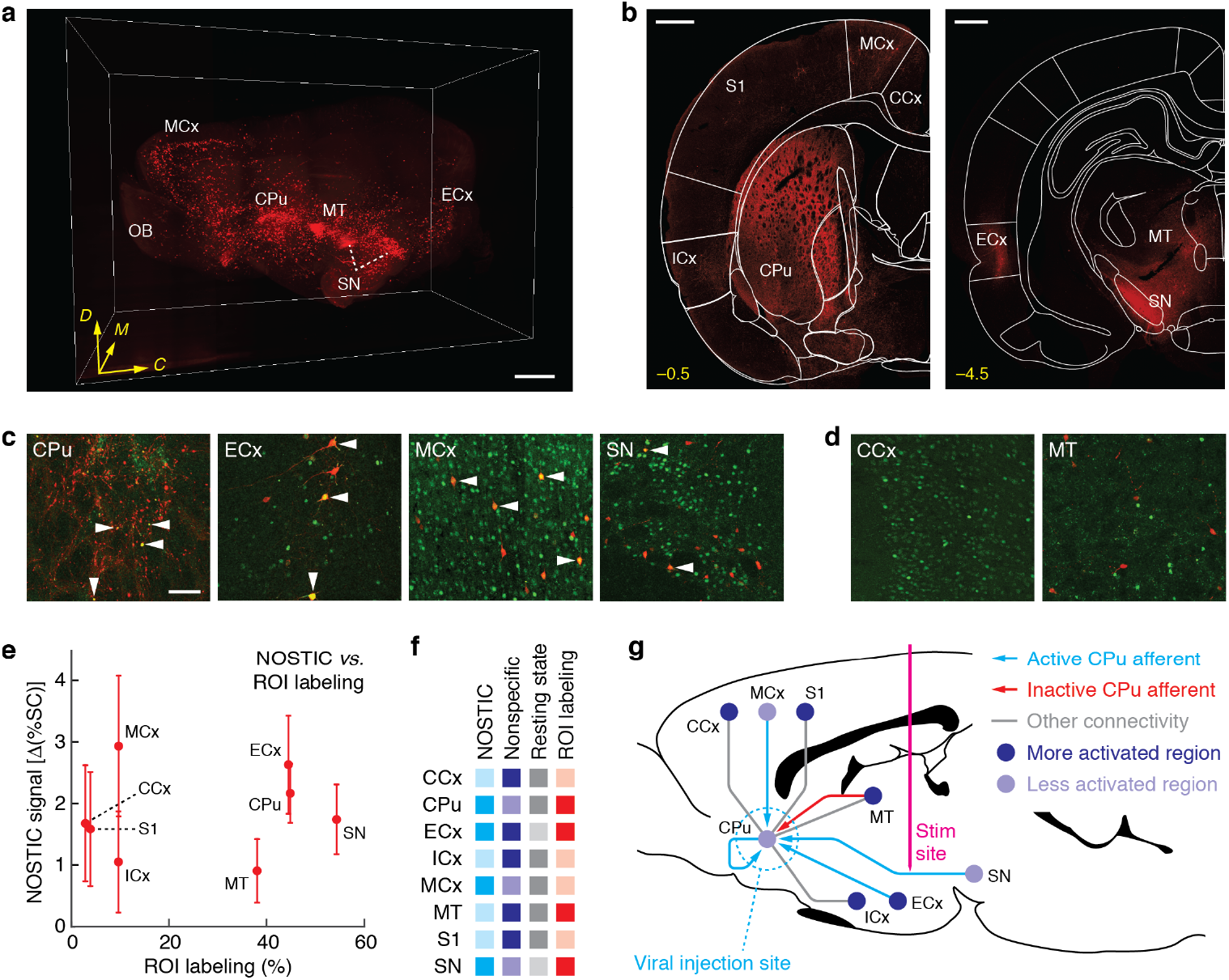
Multimodal analysis of striatal input/output relationships. **a,** Unbiased visualization of HSV-mCherry-infected brain regions in an entire cleared rat brain hemisphere following viral delivery to rat CPu. Areas with substantial expression are labeled, along with the olfactory bulb (OB), for reference. Dorsal (*D*), medial (*M*), and caudal (*C*) directions denoted in yellow. Scale bar = 2 mm. **b,** HSV-mCherry-driven expression in specific brain slices, with anatomical ROIs labeled on the atlas overlay (white) and bregma coordinates in yellow. Scale bars = 1 mm. **c,** Combined visualization of c-Fos protein (green) and HSV-directed mCherry expression (red) in animals infected with HSV-NOSTIC-1-IRES-mCherry and stimulated under conditions matching the fMRI experiments of Fig. 3. Arrowheads indicate co-stained cells in ROIs for which significant NOSTIC-mediated responses were observed. Scale bar = 100 μm. **d,** Data from two regions with-out significant NOSTIC responses, showing c-Fos staining only in CCx (top) and non-overlapping c-Fos and mCherry staining in MT (bottom). **e,** Quantification of HSV-driven expression as a percentage of stained ROIs in **b**, plotted versus NOSTIC response amplitudes. Error bars denote SEM (*n* = 6). **f,** Summary of hemogenetic imaging, nonspecific fMRI, resting state connectivity, and anatomical connectivity amplitudes across eight ROIs (left); more saturated colors denote signifi-cant NOSTIC responses (cyan), mean nonspecific fMRI responses > 3% (indigo), mean resting state *Z* values > 0.8 (gray), and anatomical labeling > 30% (red). **g,** Sagittal brain diagram sche-matizing striatal circuitry consistent with the combined results. Circles denote DBS-evoked activation levels; lines denote functional connectivity defined by hemogenetic imaging (cyan), resting state analysis (gray, non-directional), or histology (red).

The combined brain-wide dataset consisting of circuit-specific hemogenetic readouts, non-specific fMRI, resting state analysis, and anatomical tracing of striatal afferents yields an integrated view of neural information flow in and around the rodent striatum (Fig. 4f,g). The NOSTIC-1 readouts identify ECx, MCx, and SN as distal input sources that shape striatal activity during stimulation, complementing presumed local processing within CPu itself. Concurrent activation of these cortical and midbrain afferents during LH-targeted DBS has not previously been documented, but might be important for behaviorally reinforcing properties of the stimulation.^23,24^ Meanwhile, cingulate cortex (CCx), ICx, MT, and S1 are all strongly activated by LH stimulation and also correlated to CPu in the resting state, but these regions do not display statistically significant pre-synaptic input to CPu according to the NOSTIC-mediated fMRI measurements. A simple hypothesis to explain these results is that CCx, ICx, MT, and S1 are excited by striatal outputs, but we cannot rule out the possibility that alternative conduction pathways are involved; further application of hemogenetic imaging could resolve this issue. Medial thalamic input to striatum is notably not observed, despite strong anatomical and resting state connectivity between MT and CPu and activation of MT by LH stimulation. These input/output relationships might collectively generalize to other experimental contexts, particularly those related to reward processing.

## DISCUSSION

The results we present here introduce and apply a sensitive strategy for brain-wide functional imaging of genetically targeted cells and circuit elements. NOSTIC probes transduce intracellular calcium signaling into hemodynamic responses that can be measured using fMRI and other non-invasive neuroimaging modalities. Our studies demonstrate efficacy of the probes and their suitability for selective fMRI of neural populations labeled using a viral vector. By combining NOSTIC-based hemogenetic imaging with nonspecific fMRI and postmortem histological analysis, we define distinct relationships between function and anatomy in striatal circuitry during reward-related DBS.

Our analysis of striatal functional connectivity using the NOSTIC-1 activity reporter has broader importance in several respects. First, our results demonstrate the complexity of responses to DBS. Although the LH stimulation we used is targeted at fibers that directly connect midbrain nuclei to CPu, our results demonstrate engagement of neural pathways that extend far beyond these structures. The activation of distal CPu afferents from cortex we observed could not be anticipated based on anatomical considerations alone, and yet might be critical for behavioral consequences of the stimulation paradigm. Our data thus imply that both direct and indirect effects of targeted stimulation are important in determining how DBS affects the brain.

Second, our results indicate limitations of conventional approaches for understanding brain-wide connectivity. Anatomical tracing fails to predict which presynaptic inputs to the striatum are most relevant to the stimulus paradigm we explored, and nonspecific fMRI assessments during stimulation and resting states fail to highlight monosynaptic functional relationships identified by the hemogenetic imaging approach. NOSTIC-based imaging thus provides mechanistically informative brain-wide information that is unavailable using other neural activity readouts, including the combination of fMRI with cell-specific stimuli such as optogenetic manipulations.^25,26^

Finally, the molecular neuroimaging strategy we used can be extended to many other experimental contexts. In the future, hemogenetic techniques could be applied in sedated or awake animals of many species, using virtually any wide-field imaging modality, and in conjunction with spatial or genetic targeting methods designed to achieve specificity for neural or non-neural cell populations, perhaps even with intracellular precision using constructs such as NOSTIC-1P. Such approaches might be particularly powerful for analyzing the basis of functional selectivity in high level sensory processing, or for characterizing the roles of distinct cell types and projections in neural mechanisms of behavior. Further protein engineering could yield NOSTICs that function under conditions where endogenous hemodynamics are suppressed;^27^ this would allow identification of NOSTIC-dependent signals without the subtractive approach of Fig. 3. Similar engineering could yield NOSTICs suitable for multiplexed measurements from more than one cell population in the same animal. Variants of the probes presented here could also be used to study processes such as neuroplasticity, neurochemical signaling, and neurovascular coupling. Hemogenetic technology thus offers a general strategy for mechanistic dissection of brain function on a previously inaccessible scale.

## Abbreviations

(BOLD): Blood Oxygen Level-Dependent
(CPu): Caudate-Putamen
(CCx): Cingulate Cortex
(DBS): Deep Brain Stimulation
(ECx): Entorhinal Cortex
(fMRI): Functional Magnetic Res-onance Imaging
(GFP): Green Fluorescent Protein
(HSV): Herpes Simplex Virus
(iNOS): Induci-ble Nitric Oxide Synthase
(ICx): Insular Cortex
(LH): Lateral Hypothalamus
(L-NAME): L-nitroargi-nine methyl ester
(MRI): Magnetic Resonance Imaging
(MT): Medial Thalamus
(MCx): Motor Cortex
(nNOS): Neuronal Nitric Oxide Synthase
(NOSTIC): Nitric Oxide Synthase for Targeting Image Contrast
(S1): Primary Somatosensory Cortex
(SN): Substantia Nigra

## Author Contributions

SG, NL, BBB, and AJ designed the research. SG, NL, BBB, TX, JD, and UDS performed the *in vitro* and *in vivo* experiments. KX, ND, NDE, and KC implemented the brain clearing and associated histology procedures. SG, NL, and AJ analyzed the results and wrote the paper.

## Acknowledgements

This research was funded by NIH grants R01 DA038642, R24 MH109081, UF1 NS107712, and U01 NS103470 and a grant from the MIT Simons Center for the Social Brain to AJ. SG was supported by an HHMI International Student Research Fellowship and Sheldon Razin Fellowship from the McGovern Institute for Brain Research. NL was supported by a Stanley Fahn Research Fellowship from the Parkinson’s Disease Foundation. TX was a visiting student from the Beijing University of Chinese Medicine, funded by a scholarship from the China Scholarship Council. JD was supported by the Johnson & Johnson UROP Scholars Program at MIT.

The authors are grateful to Sabbi Lall and Bernardo Sabatini for detailed comments on the manuscript. They also thank Thomas Poulos of the University of California, Irvine and Paul Ortiz de Montellano at the University of California, San Francisco for providing NOS constructs. Rachael Neve of the Massachusetts General Hospital is acknowledged for production of HSV vectors.

## METHODS

### Animal subjects

Adult male Sprague-Dawley rats (200-250 g) were purchased from Charles River Laboratories (Wilmington, MA). After arrival, animals were housed and maintained on a 12 hr light/dark cycle and permitted *ad libitum* access to food and water. All procedures were carried out in strict compliance with National Institutes of Health guidelines, under oversight of the Committee on Animal Care at the Massachusetts Institute of Technology.

### Construction of engineered NOS variants

Plasmids encoding NOS constructs were a generous gift from the lab of Dr. Thomas Poulos at the University of California, Irvine. The nNOS and iNOS genes were individually cloned into the pIRESpuro3 mammalian vector (Takara Bio USA, Mountain View, CA). An inverse polymerase chain reaction (PCR) was performed to create the backbone. Inserts were amplified using overhangs ranging from 24-30 nucleotides on each end. A single-step, isothermal, ligation-free procedure was used to integrate the inserts. The entire length of the coding strand was verified using sequencing. A similar DNA assembly strategy was used to generate chimeric genes encoding NOSTIC-1 and NOSTIC-1P.

The following strategy was used to build genetic constructs for making stable cell lines: A lentiviral backbone containing the cytomegalovirus (CMV) promoter driving enhanced green fluorescent protein (GFP) with a blasticidin resistance cassette driven by a phosphoglycerate kinase (PGK) promoter (pLV6-eGFP-BLA) was used to generate pLV6-NOSTIC-1-BLA. The parent plasmid was treated with BamHI (NEB, Ipswich, MA) and EcoRI (NEB, Ipswich, MA) to make linearized vector. NOSTIC-1 was amplified using PCR with primers having 20-25 bp overhang. The purified PCR product and the linearized backbone were assembled using Gibson assembly (New England Biolabs, Ipswich, MA). DNA was transformed into Stable competent cells (New England Biolabs). The entire coding strand was verified by Sanger sequencing (Quintara Biosciences, Boston, MA).

### Expression and activity of NOS clones in cell culture

FreeStyle 293-F cells were obtained from Thermo Fisher (Waltham, MA) and maintained according to the supplier’s protocol. To test for expression and activity of NOS constructs expressed in these cells, high-purity DNA suitable for transfection was obtained using the Qiagen Plasmid Maxi Kit (Hilden, Germany). DNA was introduced into the cells using 293Fectin (Thermo Fisher) following the manufacturer’s protocols. 48 hours post-transfection, cells were pelleted and resuspended in Freestyle 293 Expression Medium (Thermo Fisher) containing 1 mM L-arginine and 1 mM CaCl_2_ and seeded in six-well plates at two million cells per well. Cells were stimulated with 5 μM A23187 (MilliporeSigma, Burlington, MA). Catalytic activity of NOS constructs was determined by quantifying nitrite from the supernatant using the Griess test (Promega, Madison, WI). For experiments to test for inhibitor sensitivity of different NOS constructs, *N*_ω_-nitro-L-arginine methyl ester hydrochloride (L-NAME) and 1400W were purchased from MilliporeSigma and added to resuspension medium prior to stimulation of the cells. 293FT cells (Thermo Fisher, Waltham, MA) were used to generate stable lines and were maintained in accordance with manufacturer’s guidelines. Cells were co-transfected with either pLV6-NOSTIC-1-BLA or pLV6-eGFP along with plasmids, psPAX2 and pMD2.G (Addgene, Watertown, MA) using Lipofectamine 3000 (Thermo Fisher, Waltham, MA). 48 hours post-transfection the supernatant containing lentiviral particles was added to freshly seeded cells on a 10-cm tissue culture dish. 24-hours following the addition of the lentiviral supernatant, the media was aspirated and changed with fresh Dulbecco’s Modified Eagle Medium (DMEM) with 10% fetal bovine serum (FBS) (Thermo Fisher) containing 10 μg/ml blasticidin. Media was changed every 48 hours until the dish was about 80% confluent.

Catalytic activity of the stably expressing NOSTIC cell line was tested using the Griess test after stimulation with A23187 as described above. To test for expression of NOSTIC, cells either stably expressing eGFP or NOSTIC-1 were seeded on poly-D-lysine coated cover slips (Thermo Fisher, Waltham, MA). When these cells were about 60% confluent, they were fixed with 4% paraformaldehyde (PFA) for 15 minutes, rinsed thrice with phosphate-buffered saline (PBS), permeabilized with buffer containing PBS and 3% TritonX (MilliporeSigma) for 15 minutes, and blocked with buffer containing PBS and 10% donkey serum (MilliporeSigma) for 60 minutes. The cells were then incubated with primary antibody (goat-anti-nNOS, Abcam, Cambridge, MA) at 1:200 dilution at 4 °C. Following overnight incubation, cells were rinsed thrice with PBS and incubated with secondary antibody (donkey anti-goat IgG, tagged with Alexa Fluor 488, Thermo Fisher) at 1:500 dilution for 1 hour. The cover slips were rinsed thrice and mounted on glass slides with ProLong Antifade Mountant (Thermo Fisher). These were then imaged on a confocal microscope (LSM 710, Carl Zeiss, Oberkochen, Germany).

### Brain implantation of NOSTIC-expressing cells

293FT cells stably expressing NOSTIC-1 or GFP were harvested by trypsinization, pelleted by centrifugation for 10 minutes at 300*g*, washed with cold PBS (pH 7.4), and resuspended at a density of 100,000 cells/μL in artificial cerebrospinal fluid (aCSF, 125 mM NaCl, 2.5 mM KCl, 1.4mM CaCl_2_, 1 mM MgCl_2_, 1.25 mM NaH_2_PO_4_, 25 mM NaHCO_3_).

Prior to intracranial cell implantation in rats, each animal was anesthetized with isoflurane (4% for induction, 2% for maintenance) and positioned in a stereotaxic instrument (Kopf Instruments, Tujunga, CA) with a water heating pad (Braintree Scientific, Braintree, MA) to keep the body temperature at 37 °C. The scalp was surgically retracted and two small holes were drilled into the skull above the target sites 0.5 mm posterior and ±3 mm lateral to bregma. 28G polyether ether ketone (PEEK) cannula guides (Plastics One, Roanoke, VA) designed to project 1 mm below the surface of the skull were lowered through the holes. The guides were affixed in place using SEcure light curing dental cement (Parkell, Edgewood, NY). A custom-fabricated plastic headpost was placed in front of the guide cannula and also secured with dental cement (Parkell).

To deliver cells, 33G metal internal cannulae (Plastics One) were lowered through the guide cannulae to a depth of 5.5 mm below the skull. Approximately 500,000 NOSTIC-expressing or control cells were infused via the cannulae at a flow rate of 0.2 μL/min for 25 min, with test and control sites alternating among experiments to negate potential biases. 10 minutes after the infusion the cannulae were slowly removed.

### Magnetic resonance imaging of implanted cells *in vivo*

After implantation of NOSTIC-expressing or control GFP-expressing cells, the metal cannulae were removed. Animals were intubated and ventilated with a small animal ventilator (Harvard Apparatus) which operated at 62 beats per minute with a 6 mL stroke volume, delivering oxygen and air as a 5:1 ratio mixture. Each animal was then transferred and fixed into a custom rat imaging cradle. A surface receive-only radiofrequency coil (Doty Scientific, Columbia, SC) was positioned over the head. 33G PEEK internal cannulae (Plastics One) were connected to PE-50 tubing (Plastics One) loaded with the calcium ionophore A23187, with or without the NOSTIC-1 inhibitor 1400W, and was lowered through the guide cannulae to the same site for cells implantation, at a depth of 5.5 mm below the skull. Isoflurane was discontinued and the animal was sedated with medetomidine (IP bolus of 1 mg/kg, followed by 1 mg/kg/hr infusion) and paralyzed with pancuronium (IP bolus of 1 mg/kg, followed by 1 mg/kg/hr infusion). Each animal was then transferred to a 9.4 T Bruker (Billerica, MA) MRI scanner for imaging. Animals were warmed using a water-heating pad (Braintree). Heart rate and blood oxygenation saturation level were continuously monitored using an MRI-compatible noninvasive infrared pulse oximeter (Nonin Medical, Plymouth, MN). Breathing rate and CO_2_ level were continuously monitored using MRI-compatible SurgiVet CO_2_ monitor (Smiths Medical, Dublin, OH).

All MRI images were obtained using a transmit-only 70 cm inner diameter linear volume coil (Bruker) and a 2 cm diameter receive-only surface coil (Doty) for excitation and detection, respectively. Scanner operation was controlled using the ParaVision 5.1 software (Bruker). High-resolution anatomical MRI images were acquired using a *T_2_*-weighted rapid acquisition with refocused echoes (RARE) pulse sequence with RARE factor = 8, bandwidth (BW) =200 kHz, effective echo time (*TE*) = 30 ms, repetition time (*TR*) = 5 s, in-plane field of view (FOV) = 2.56 × 1.28 cm, in-plane resolution of 100 μm × 100 μm, and slice thickness = 1 mm. Functional imaging scan series were acquired using a multi-echo echo-planar imaging (ME-EPI) pulse sequence *TE* values = 10, 23, and 36 ms, flip angle (FA) = 90°, *TR* = 2 s, FOV = 2.56 × 1.28 cm, and in-plane resolution of 400 × 400 μm over five coronal slices with slice thickness = 1 mm.

MRI scans were continuously collected for 2 minutes baseline, 5 minutes during calcium stimulation or control treatment, and 10 minutes resting after the stimulation. Calcium stimulation was produced by infusion of a solution of A23187 in aCSF, at a rate of 0.2 μL/min through the preimplanted plastic cannulae. Control treatment involved infusion of an identical solution that also contained 1400W. *T_2_\**-dependent contributions to the functional imaging data were extracted from the multi-echo image time series by examining data at three echo times, permitting voxels with strong *TE* dependence characteristic of BOLD contrast to be identified.^28^ Non-*T_2_\**-weighted signal components, largely along the cell implantation tracks, were thus removed from the analysis. The preprocessed data were then further analyzed in MATLAB (Mathworks, Natick, MA) for visualization and quantification of results. Response maps were computed as mean percent signal change during the expected peak response time point 1-100 s after A23187 stimulation onset minus the average baseline signal, 1-100 s before stimulus onset. Mean response amplitudes and time courses were evaluated over an ROI defined around each cell implantation site. These ROIs were defined by the edges of the implanted xenografts, as judged from the high-resolution *T_2_*-weighted anatomical scans, expanded by four voxels in each in-plane direction.

### Preparation of herpes simplex viral vectors

Herpes Simplex Virus-1 (HSV) vectors used in this study were produced by Rachael Neve at the Massachusetts General Hospital Gene Delivery Technology Core. The vectors are replication deficient and capable of retrograde transport from injection sites in rodent brain. NOSTIC-1 expression was directed by an HSV containing the NOSTIC-1 gene in tandem with the gene for mCherry, separated by an internal ribosome entry site (IRES), with expression directed by the human EF1α promoter (HSV-hEF1α-NOSTIC-1-IRES-mCherry). A control vector directed expression of mCherry only from the same promoter (HSV-hEF1a-mCherry).

Virally-delivered NOSTIC-1 was tested for catalytic activity in cell culture. Approximately 200,000 HEK 293FT (Thermo Fisher) cells were seeded on poly-D-lysine and laminin coated coverslips (Thermo Fisher) in a 24-well plate. The following day, 1 μL of 10^9^ pfu/mL HSV delivering NOSTIC-1 or mCherry was added to each well. 24-48 hours later, media was aspirated, washed, and the cells were stimulated with 5 μM A23187 in 400 μL FreeStyle 293 Expression medium (Thermo Fisher) per well, supplemented with 1 mM CaCl_2._ Nitrite was quantified from supernatants using the Griess assay following overnight incubation.

### Stereotaxic injection of viral vectors

Animals were prepared for cranial surgery to expose the desired viral injection site at the anterior central caudate-putamen (CPu). Each animal was anesthetized with isoflurane (4% for induction, 2% for maintenance). A small hole was drilled into the skull above the target site, 0.5 mm posterior and 3 mm lateral to bregma. 35G metal internal canulae connected with Hamilton syringes preloaded with each viral vector (NOSTIC-1: 1 × 10^9^ pfu/mL, mCherry: 2 × 10^9^ pfu/mL) were then lowered into the brain and each vector was infused at a rate of 0.1 μL/min at two sites (3 μL per site), 6 mm and 5 mm below the skull surface. The canulae were slowly removed 10 minutes after the viral injection. Bone wax was applied to each injection site to seal it using a sterile cotton applicator tip. Skin incisions were closed by sterile wound clips and lidocaine gel (2%) was applied over the wound areas. Isoflurane was then discontinued and each rat was removed from the stereotaxic frame and placed in warmed cage on a heating pad to recover for 45 min. Slow-release buprenorphine (0.3 mg/kg, MIT Pharmacy) was administered subcutaneously to minimize pain and discomfort. Wound clips were removed 7-10 days after surgery and when the wounds had healed.

### Implantation of stimulation electrodes

Three weeks after viral injection, rats underwent a second surgery to implant electrodes suitable for deep brain stimulation (DBS) of the medial forebrain bundle region of the lateral hypothalamus (LH). Animals were prepared similarly for cranial surgery to expose the desired stimulation site. Bipolar stimulating electrodes were targeted to coordinates 3.4 mm posterior, 1.7 mm medial, and 8.6 mm ventral to bregma. In addition, a plastic head-post was placed to facilitate positioning of each animal during MRI scanning. After all the items were secured on the skull, dental cement was applied to the entire exposed skull surface area to hold them rigidly in place. After the surgery, animals were maintained under a heating pad and treated with 0.3 mg/kg slow-release buprenophine subcutaneously during a convalescence period. Rats were allowed 4-7 days for recovery prior to fMRI experiments.

### Functional magnetic resonance imaging

Preparation of animals for fMRI was similar to that described above for xenograft implantation experiments. After intubation via a tracheotomy and ventilation, each animal was transferred to the MRI cradle and fixed by the headpost. For imaging of rats during LH stimulation, each animal’s LH electrode was connected to a constant-current stimulus isolator (World Precision Instruments, Sarasota, FL) controlled from a laptop computer. For imaging of rats during somatosensory stimulation, each animal was implanted with subcutaneous electrodes in the forelimb, which were similarly connected to the stimulus isolator.

General MRI methods, including hardware and anatomical imaging, were equivalent to those described above for xenograft-implanted animals. Functional imaging of HSV-transfected rats included two minutes of baseline image acquisition followed by five cycles of electrical stimulation, each consisting of 16 s LH stimulation followed by 5 min of rest. Each 16 s stimulation block consisted of eight 1 s trains of 1 ms 0.17 mA current pulses delivered at a frequency of 60 Hz, separated by 1 s between trains. Functional imaging with forepaw stimulation consisted of six blocks of 3 mA 1 ms pulses repeated at 9 Hz for 10 s alternating with 40 s of rest. In each case, the stimulation paradigm was synchronized with the image acquisition using a custom script exe-cuted in LabVIEW (National Instruments, Austin, TX). To minimize potential radiofrequency artifacts, the stimulator cable was filtered with a 60 MHz low-pass filter (Mini-Circuits, Brooklyn, NY) before entering the MRI enclosure.

A *T_2_\**-weighted echo planar imaging (EPI) sequence was used for detection of stimulus-induced BOLD contrast, with BW = 200 kHz, *TE* = 16 ms, FA = 90, *TR* = 2s, FOV = 2.56 × 1.28 cm, in-plane resolution 400 × 400 μm, and slice thickness = 1 mm. In order to discern NOS inhibitor dependence of fMRI signals, image series from each animal were acquired first in the absence and then in the presence of 1400W or L-NAME. For hemogenetic experiments, identical procedures were applied using animals infected with NOSTIC-encoding and control viruses.

### Analysis of functional imaging data

Images were reconstructed using the ParaVision 5.1 software and then imported into the National Institute of Health AFNI software package^29^ for further processing. High-resolution anatomical images of each animal were registered to a Waxholm co-ordinate space rat brain atlas.^30^ Functional imaging time series were preprocessed in steps that included slicetiming correction, motion correction using a least-squares rigid-body volume registration algorithm, voxelwise intensity normalization, spatial smoothing with Gaussian spatial kernel of 0.5 mm full width at half maximum, and spatial resampling to double the image matrix size. Segmentation of brain from non-brain voxels was performed in MATLAB. Preprocessed time series were then co-registered onto the previously atlas-aligned anatomical images. Regions of interest (ROIs) used for subsequent analyses were also defined with respect to the rat atlas and are displayed in Supplementary Fig. 1.

For image analysis of HSV-transfected rats, stimulus-dependent *T_2_\**-weighted EPI time courses were estimated using the 3dDeconvolve general linear modeling implementation in AFNI. LH stimulus time blocks were used as event regressors. Six motion correction parameters from each animal were included as nuisance regressors, along with a linear baseline term. Outlier scans detected by median deviation from time series trends in each data set were censored from the analysis. Mean stimulus response functions of 242 s duration (121 image frames) were obtained in units of percent signal change for each voxel in each animal.

Mean percent signal change maps were computed as the mean signal for 174 s (87 image frames) beginning at stimulus onset, minus the baseline signal defined by the mean of response at time points 0-40 s (20 frames) before stimulus onset. To correct for systematic changes in physiology between pre-and post-1400W conditions, fMRI responses obtained after 1400W treatment were scaled so that the hemodynamic response along the midline (assumed to be NOSTIC-independent) matched that observed before treatment. Difference maps were then calculated by subtracting the mean response map with 1400W treatment from the corrected mean response map without 1400W treatment from the same set of animals. ROI-specific values were obtained by averaging fMRI signals over the corresponding regions. Unless otherwise noted, SEM values associated with mean amplitudes, time courses, and other response parameters were computed by jackknife resampling over multiple animals.

For resting state connectivity analysis, mean signal time courses for each ROI were calculated from EPI image series acquired from four naïve rats in the absence of stimulation. The Pearson correlation coefficients and the corresponding Fisher-transformed *Z* values were computed between the time courses of the viral injection subregion of CPu (bregma –0.5 mm) and the eight ROIs. Within-subject Student’s *t*-tests were used to evaluate the statistical significance for the correlation analysis. Bonferroni correction was applied to assess significance for multiple comparisons (*p* < 0.05/8 = 0.006).

### Brain clearing and imaging

Three weeks after the HSV-mCherry virus injection, rats were transcardially perfused with 200 mL of ice-cold phosphate-buffered saline containing 0.02% sodium azide (PBSN), followed by 200 mL of ice-cold SHIELD perfusion solution at a flow rate of 60 mL/min.^22^ The brain was extracted and incubated in SHIELD perfusion solution at 4 °C on an orbital shaker for 2 days. The brain was then moved to 50 mL of SHIELD OFF solution and incubated on an orbital shaker for 3 days at 4°C. After SHIELD OFF incubation, the brain was incubated in 40 mL of SHIELD ON solution that had been pre-warmed at 37 °C for 20 minutes. SHIELD ON incubation occurred on an orbital shaker at 37 °C for 1 day. Following the completion of SHIELD processing, the brain was washed in PBSN for at least 1 day. SHIELD processing solutions were obtained from LifeCanvas Technologies (Cambridge, MA).

The brain was hemisected after washing. The meninges surrounding the brain were then stained using a solution of 1% toluidine blue in 70% ethanol, diluted 10-fold in PBSN. The brain was briefly immersed in the staining solution and then placed under a stereoscope equipped with a fiber-optical illuminator. A blunt-tipped paint brush was used to separate the dura layer from the surface of the brain. Hemispheres were then passively incubated for 48 hours at room-temperature in a tissue clearing buffer containing sodium dodecyl sulfate. After incubation, intact hemispheres were rapidly delipidated via stochastic electrotransport at 45 °C for 10-14 days.^31^ Optical clearing proceeded until the core of each sample showed no signs of opacity. Samples were then rinsed in PBSN three times for a minimum of 12 hours each. Prior to volumetric labeling, samples were passively incubated for 48 hours at room temperature in eFLASH sample buffer.^32^ Rapid 24 hour immunostaining of whole hemispheres with polyclonal CF640R rabbit anti-RFP (Biotium, Fremont, CA) was carried out using the eFLASH protocol.^32^ The samples were again passively washed in PBSN three times for a minimum of 12 hours each before refractive index matching.

After clearing and labelling, the intact rat hemisphere was optically cleared using a Protos-based immersion medium. The tissue was incubated for two days at 37 °C until tissue was transparent. Once optically cleared, the tissue was mounted in an agarose block (1.5% wt/vol agarose powder dissolved in immersion medium) and volumetrically imaged with an axially-swept light-sheet microscope (LifeCanvas Technologies) using a custom 3.6x magnification, 0.2 numerical aperture detection objective with a uniform axial resolution of approximately 4 μm. This system is equipped with an Oxxius (Lannion, France) L4Cc laser combiner unit equipped with 488 nm, 561 nm, 642 nm, and 785nm diodes; anti-RFP images were acquired using the 642 nm unit. The 900 GB raw light sheet dataset was stitched into a single 92 GB Imaris (Bitplane, Zürich, Switzerland) dataset for visualization.

### Conventional immunohistochemical analysis

For immunohistochemistry analysis of activity-dependent gene induction in conjunction with other markers, rats injected with HSV-hEF1a-NOSTIC-1-IRES-mCherry were anesthetized via an intraperitoneal injection of a ketamine (90 mg/kg) and xylazine (10 mg/kg) mixture. Thirty minutes after the induction of anesthesia, the rats were subjected to a sequence of five trials of LH stimulation using the same stimulation parameters as in the fMRI experiments, with 1 min rests between trials. Rats were then transferred back to their own cage for 90 minutes to allow for c-Fos expression and accumulation before the animals were sacrificed by lethal intraperitoneal injection of a sodium pentobarbital (0.5 mg/g Fatal-Plus), in conjunction with transcardial perfusion with 4% paraformaldehyde (PFA).

Brains were harvested and fixed in PFA for an additional 48 hours at 4 °C. Extracted brains were then sectioned into 50 μm-thick free-floating coronal slices using a semi-automatic vibratome (VT1200, Leica Biosystems, Wetzlar, Germany). Slices were stored in PBS at 4 °C. Prior to staining, slices were washed thrice in PBS + 0.1% TritonX (PBST) and blocked for 1 hour with 10% v/v donkey serum (MilliporeSigma). Each slice was then incubated overnight at 4 °C with gentle shaking in PBST with 1% serum and primary antibody. The following primary antibodies were used: goat anti-mCherry (1:200, Lifespan Biosciences, Seattle, WA), rabbit anti-mCherry (1:200, Abcam), rabbit anti-c-Fos (1:500, Synaptic Systems, Goettingen, Germany), goat anti-IBA-1 (1:500, Abcam), rabbit anti-nitrotyrosine (1:200, MilliporeSigma).

The following day, cells were washed thrice with PBST and then probed with secondary antibody (Thermo Fisher or Abcam). The following secondary antibodies were used: donkey antirabbit IgG tagged with Alexa Fluor 594, donkey anti-goat IgG tagged with Alexa Fluor 488, donkey anti-goat IgG tagged with Alexa Fluor 594, or donkey anti-rabbit IgG tagged with Alexa Fluor 488, all applied in PBST with 1% serum for one hour. Slices were then washed thrice in PBS, mounted on glass slides using Prolong Antifade Mountant with DAPI (Thermo Fisher) and imaged on a confocal microscope (LSM 710, Carl Zeiss, Oberkochen, Germany).

---

**Supplementary Figure 1.**
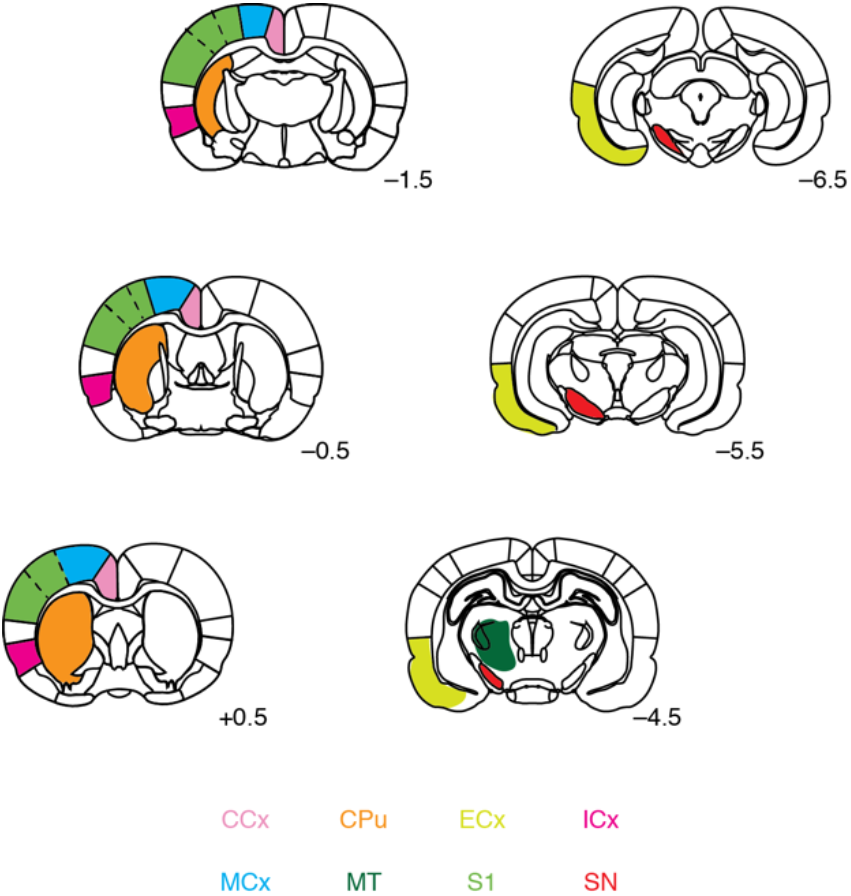
Regions of interest used in fMRI analyses. Relevant regions of interest were defined with respect to a standard brain atlas. Color coding corresponds to labels at bottom: caudate-putamen (CPu), cingulate cortex (CCx), insular cortex (ICx), motor cortex (MCx), primary somatosensory cortex (S1), substantia nigra (SN), entorhinal cortex (ECx), and central medial thalamus (CMT). Rostrocaudal coordinates relatively to bregma indicated for each coronal slice shown.

**Supplementary Figure 2.**
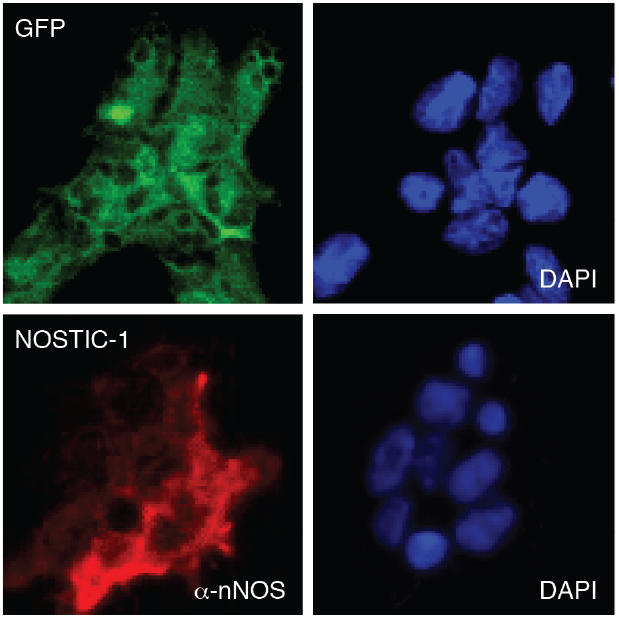
Stable expression of NOSTIC-1 and GFP in transfected cells. Stable transfects were assayed to determine expression of their respective constructs prior to imaging in implanted rat brains. GFP-transfected control cells (top) were visualized by green fluorescence and DAPI staining (right). NOSTIC-1-transfected cells (bottom) were stained with anti-nNOS antibody (α-nNOS, left) and DAPI (right).

**Supplementary Figure 3.**
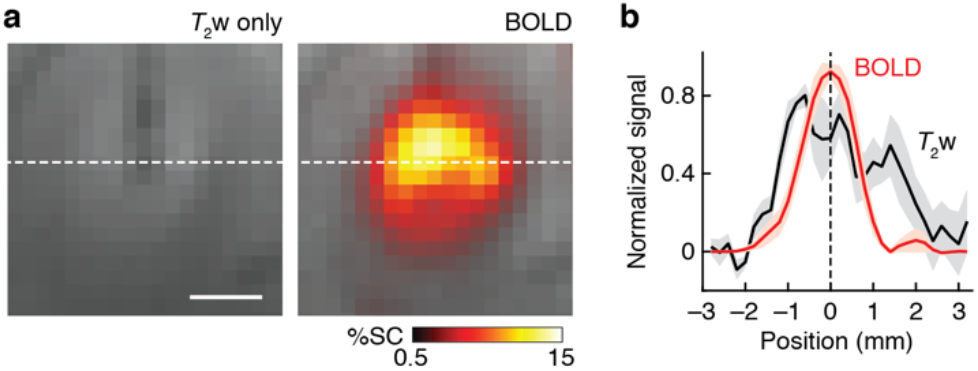
Spatial extent of xenograft-induced hemogenetic responses. **a,** Left: Anatomical MRI showing *T*_2_-weighted (*T*_2_w) contrast in the neighborhood of cell implantation for a representative animal from the experiments of main text Fig. 2. Right: Average BOLD contrast map (color) induced by A23187 stimulation in units of percent signal change (%SC), overlayed on a *T*_2_-weighted scan. **b,** Cross sections of the contrast patterns through the dotted line in panel **a**, with solid lines denoting means and shading denoting SEM across six animals’ BOLD (red) and *T*_2_w (black) signal changes. Full width at half maximum is 1.4 ± 0.1 mm for the hemogenetic BOLD responses, indicating point-spread function comparable to or less than conventional hemodynamic fMRI.

**Supplementary Figure 4.**
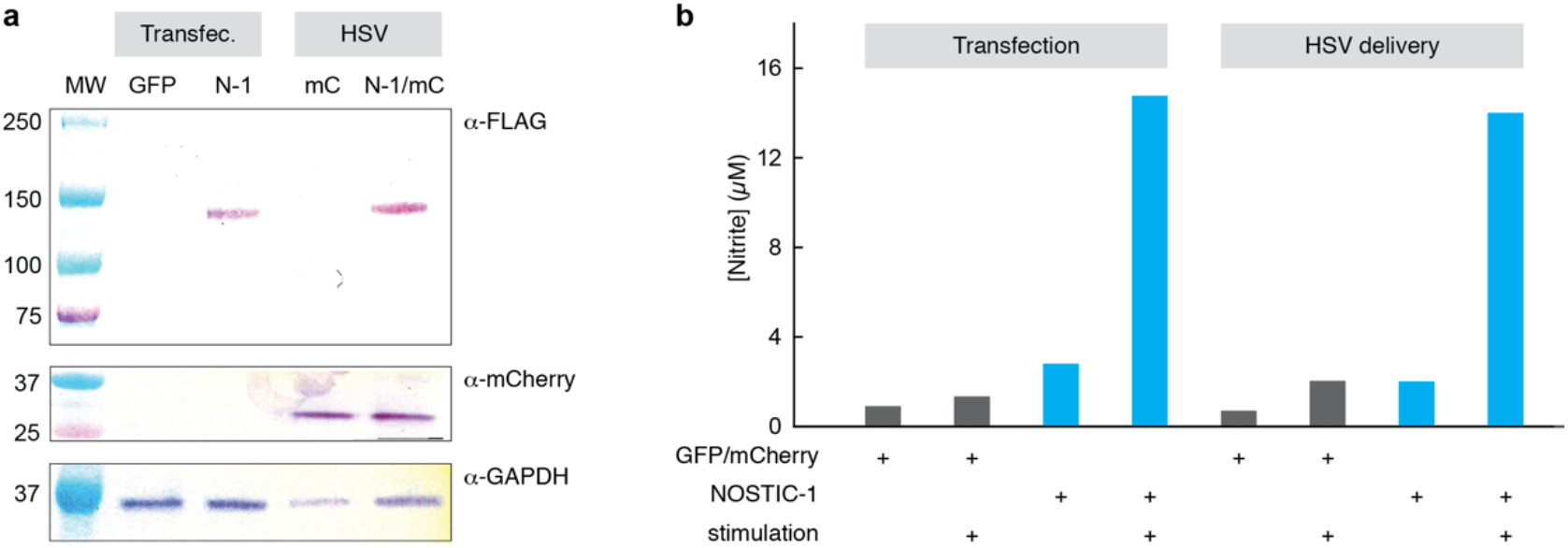
Evaluation of HSV vectors in cell culture. **a,** Western blotting was used to analyze cells transfected with GFP or NOSTIC-1 (N-1), along with cells infected with HSV encoding mCherry (mC) or NOSTIC-1-IRES-mCherry (N-1/mC). Expression and size of NOSTIC-1 was confirmed using blotting with an anti-FLAG tag (α-GAPDH) was used as a loading control (bottom). Relevant molecular weight (MW) markers labeled at left. Expected sizes: NOSTIC-1 140 kD, mCherry 29 kD, GAPDH 37 kD. **b,** Griess test measurement of nitrite production in the presence of cells transfected with GFP or NOSTIC-1 or virally transduced with mCherry or NOSTIC-1, in the presence or absence of A23187 stimulation, as indicated. Only stimulated cells expressing NOSTIC-1 show strong evidence of stimulus-dependent NO production, as indicated by the Griess test results.

**Supplementary Figure 5.**
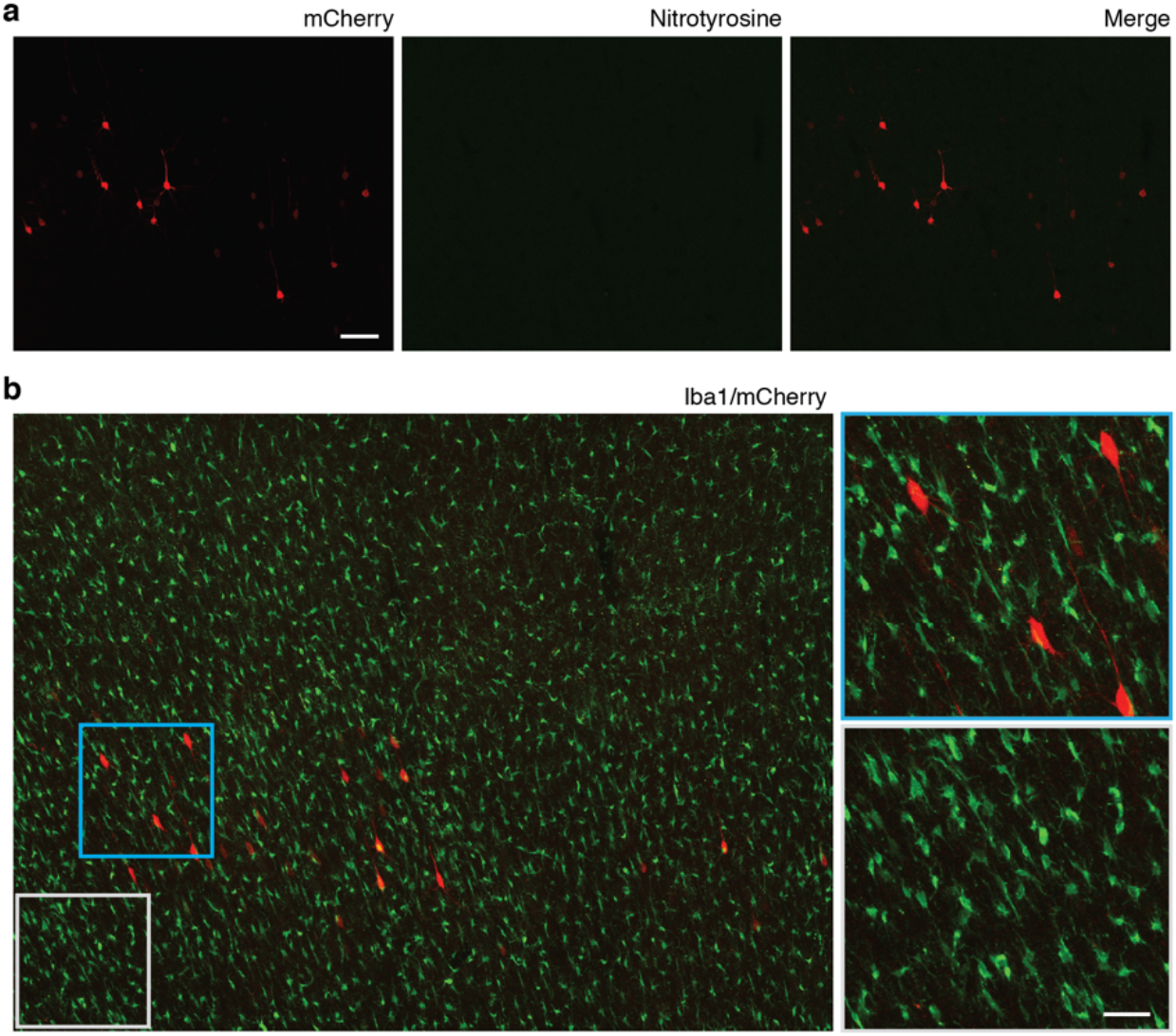
Evaluation of inflammation signatures in NOSTIC-expressing rats. **a,** Micrographs stained for mCherry (left) and nitrotyrosine (a product of excessive NO signaling, middle) show no evidence of nitrotyrosine staining in the neighborhood of cells infected with HSV-hEF1α-NOSTIC-1-IRES-mCherry in MCx. Scale bar = 100 μm. **b,** A field of cells stained for the microglial marker Iba1 (green) and mCherry (red) show no evidence of microglial activation or abnormal microglial morphology in the neighborhood of NOSTIC-encoding HSV transduction. Closeups (right) correspond to color-coded boxes (left). Scale bar = 40 μm.

**Supplementary Figure 6.**
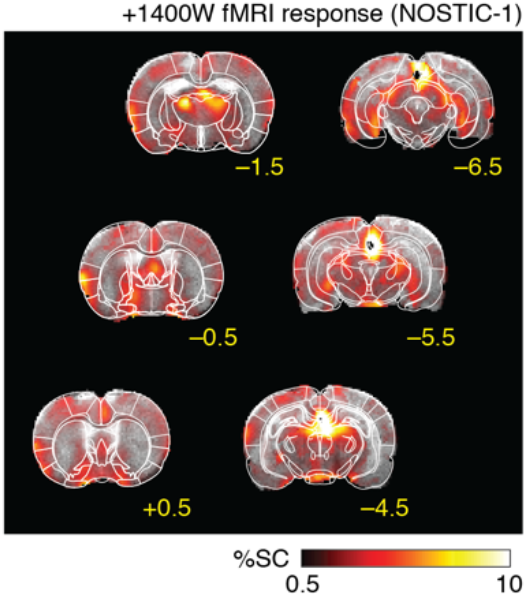
LH stimulus responses in NOSTIC-expressing rats after 1400W treatment. Average fMRI responses to LH stimulation in the +1400W condition, among 6 animals infected with NOSTIC-encoding HSV, analogous to −1400W condition data in main text Fig. 1B. Significant responses with *F*-test *p* ≤ 0.01 shown.

**Supplementary Figure 7.**
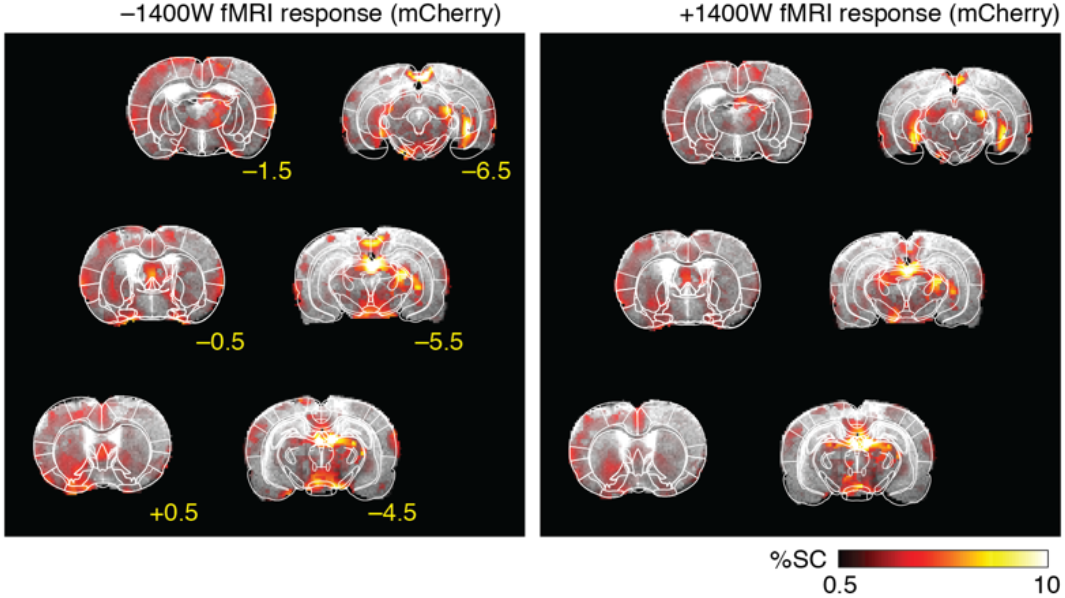
LH stimulus responses in rats treated with control HSV vectors. Average fMRI responses to LH stimulation in the −1400W (left) and +1400W (right) conditions, among 5 animals infected with mCherry-encoding HSV, corresponding to difference maps shown in main text Fig. 3C. Significant responses with *F*-test *p* ≤ 0.01 shown.

